# Structure and genome ejection mechanism of *Podoviridae* phage P68 infecting *Staphylococcus aureus*

**DOI:** 10.1101/447052

**Authors:** Dominik Hrebík, Dana Štveráková, Karel Škubník, Tibor Füzik, Roman Pantůček, Pavel Plevka

**Affiliations:** Central European Institute of Technology, Masaryk University, Kamenice 5, 625 00, Brno, Czech Republic; Faculty of Science, Masaryk University, Kamenice 5, 625 00, Brno, Czech Republic

**Keywords:** structure, *Staphylococcus aureus*, bacteriophage, cryo-electron microscopy

## Abstract

Phages infecting *S. aureus* have the potential to be used as therapeutics against antibiotic-resistant bacterial infections. However, there is limited information about the mechanism of genome delivery of phages that infect Gram-positive bacteria. Here we present the structures of *S. aureus* phage P68 in its native form, genome ejection intermediate, and empty particle. The P68 head contains seventy-two subunits of inner core protein, fifteen of which bind to and alter the structure of adjacent major capsid proteins and thus specify attachment sites for head fibers. Unlike in the previously studied phages, the head fibers of P68 enable its virion to position itself at the cell surface for genome delivery. P68 genome ejection is triggered by disruption of the interaction of one of the portal protein subunits with phage DNA. The inner core proteins are released together with the DNA and enable the translocation of phage genome across the bacterial membrane into the cytoplasm.

## Introduction

Phages from the family *Podoviridae* have complex virions composed of genome-containing heads and short non-contractile tails. The tail is attached to a special fivefold vertex of the head in which a pentamer of capsid proteins is replaced by a portal complex. The tails of podoviruses are formed of lower collar proteins, knobs, spikes, and fibers ^1^. Podoviruses that infect *S. aureus* often use cell wall teichoic acid as a receptor^2,3^. After binding to a cell, podoviruses disrupt the bacterial cell wall and eject their genomes into the cell cytoplasm^1^.

Phages infecting *S. aureus* are of interest because of their potential use in phage therapy against antibiotic-resistant infections ^4^. *S. aureus* causes a range of illnesses from minor skin infections to life-threatening diseases such as pneumonia, meningitis, and sepsis ^5–7^. Many *S. aureus* strains, particularly those found in hospitals, carry antibiotic resistance genes ^8^,^9^. Annual medical expenses caused by *S. aureus* in the United States and European Union have been estimated to exceed $2.5 billion ^10–12^.

Bacteriophage P68 belongs to the subfamily *Picovirinae*, and the genus *P68virus* of phages infecting *S. aureus*^13^. P68 has a 18,227-bp-long double-stranded DNA genome that encodes 22 open reading frames^13^. Here we used a combination of cryo-electron microscopy (cryo-EM) and X-ray crystallography to structurally characterize the virion of phage P68 and the mechanism of its genome ejection.

## Results and Discussion

### Virion structure of phage P68

The virion of P68 has an icosahedral head with a diameter of 480 Å and 395 Å-long tail, which is decorated with tail fibers (Fig. 1ab, Table S1, S2). Electron micrographs of a purified P68 sample contained not only native virions but also particles that were in the process of genome release and empty particles (SFig. 1). The complete structure of the native virion of P68 was determined to a resolution of 4.7 Å (SFig. 2, 3, Table S1). The structures of the capsid and tail with imposed icosahedral and twelvefold symmetries, respectively, were determined to resolutions of 3.3 and 3.9 Å (SFig. 2, 3, 4, Table S1).

**Fig. 1.**
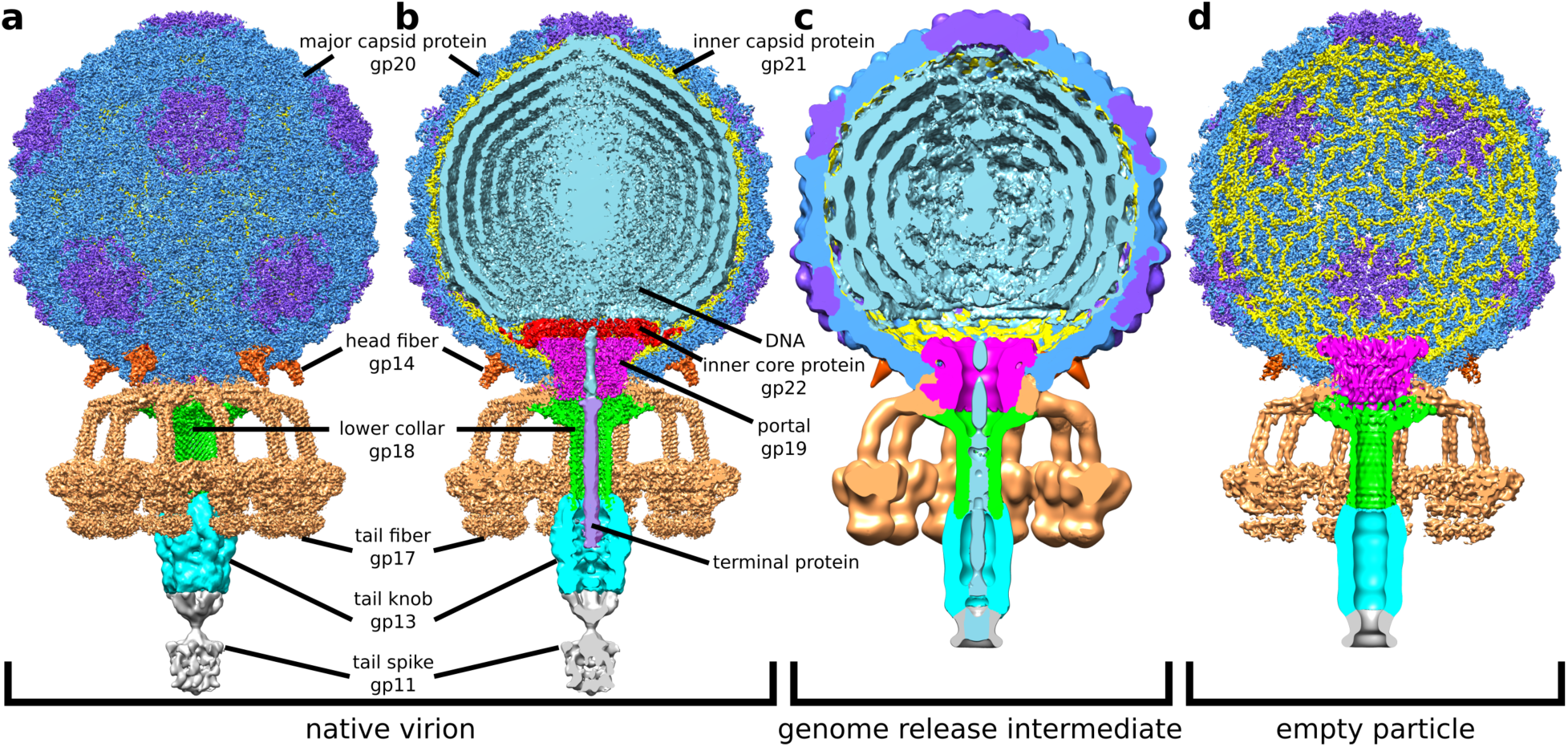
Structures of P68 virion (a, b), genome release intermediate (c), and empty particle (d). The whole P68 virion is shown in (a), whereas particles without the front half are shown in (b-d). The structures are colored to distinguish individual types of structural proteins and DNA.

### P68 capsid structure

The capsid proteins in the P68 head are organized in a T = 4 icosahedral lattice (Fig. 2ab) ^14^. The major capsid protein gp20 has the canonical HK97 fold common to numerous tailed phages and herpesviruses ^15–17^. According to the HK97 convention, the protein can be divided into four domains: the N-terminal arm (residues 1-84), extended loop (85-124), peripheral domain (125-182 and 346-388), and axial domain (183-277, 341-345, and 389-408) (Fig. 2c). Unlike in HK97, the P68 major capsid protein also contains an insertion domain (residues 278-340). The insertion and peripheral domains form a cleft that binds the extended loop of an adjacent major capsid protein and thus contribute to the capsid’s stability (SFig. 5).

**Fig. 2.**
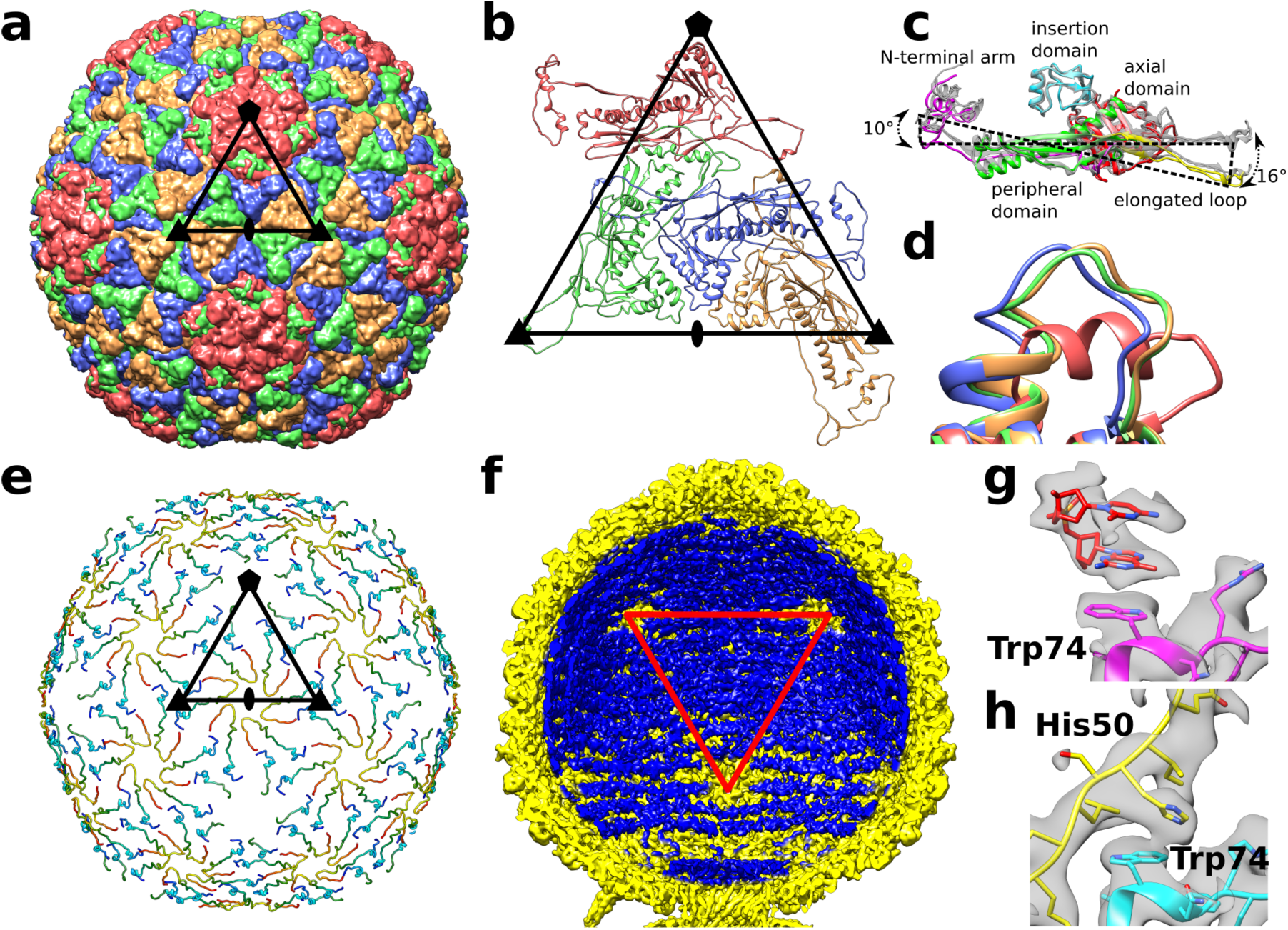
Capsid structure of P68. Major capsid proteins of P68 have HK97 fold and form T = 4 icosahedral lattice (a). Positions of selected icosahedral five, three, and twofold symmetry axes are indicated by pentagon, triangles, and oval. Borders of one icosahedral asymmetric unit are highlighted. Cartoon representation of P68 major capsid proteins in icosahedral asymmetric unit. Positions of icosahedral symmetry axes and borders of icosahedral asymmetric unit are shown. Major capsid proteins from icosahedral asymmetric unit differ in positions of elongated loops and N-terminal domains (c). Color-coding of one of the subunits indicates division of major capsid protein to domains. Residues 253-263 from the axial domain of major capsid proteins differ in structure (d). The residues form an α helix in the subunit that is part of the pentamers, whereas they constitute loops in the other subunits. The color-coding of subunits is the same as in (b). The inner capsid protein is organized in a T=4 icosahedral lattice. Proteins are rainbow colored from N-terminus in blue to C-terminus in red. Subunits related by icosahedral threefold axes and quasi-threefold axes of the T=4 lattice form three-pointed stars in which the C-terminus of one subunit is positioned next to the N-terminus of another subunit (e). Borders of a selected icosahedral asymmetric unit are shown. The ordering of the packaged P68 double stranded DNA genome (shown in blue) is disrupted around the fivefold vertices of the capsid (shown in yellow) (f). Red triangle indicates one face of icosahedron. Stacking interactions of two nucleotides with side-chain of trp 74 of major capsid protein located next to fivefold vertex (g). Side chains of trp 74 of major capsid proteins that form hexamers bind to his 50 of inner capsid proteins (h).

The quasi-equivalent structure of the T = 4 icosahedral capsid includes conformational differences in the major capsid proteins from the icosahedral asymmetric unit (Fig. 2bc). The N-terminal arms and extended loops are in one plane in the major capsid proteins that connect two hexagons (Fig. 2c). In contrast, the same domains are bent 16° in the major capsid proteins that form pentagons, and 8° in subunits that connect hexagons to pentagons (Fig. 2c). Additional differences among the capsid proteins are in the structures of residues 253-263 from the axial domain, which fold into α-helices in subunits that form pentamers and loops in subunits that belong to hexamers (Fig. 2d).

### Inner capsid proteins mediate contacts of capsid with genome

The inner face of the P68 capsid is lined by inner capsid proteins gp21, organized with icosahedral T = 4 symmetry (Fig. 2e). Except for an α-helix formed by residues 14-21, the 55-residue-long inner capsid protein “Arstotzka” lacks secondary structure elements. The inner capsid proteins related by icosahedral threefold axes and quasi-threefold axes form three-pointed stars (Fig. 2e) and are arranged so that the N-terminus of one subunit is located close to the C-terminus of another one within the stars (Fig. 2e). In contrast to protein P30 of phage PRD1 ^18^, the inner capsid proteins of P68 have limited contacts with each other.

The electron density of P68 double-stranded DNA is resolved inside the regions of the capsid lined by the inner capsid proteins, but it is missing in the proximity of fivefold vertices, where the inner capsid proteins are not structured (Fig. 2ef). The electron densities of two nucleotides of single-stranded DNA are stacked against the side chains of trp 74 of the major capsid proteins that form pentamers (Fig. 2g). In contrast, the side chains of trp 74 of major capsid proteins that form hexamers interact with the side chains of his 51 of the inner capsid proteins (Fig. 2h), and thus cannot bind phage DNA. Therefore, the inner capsid proteins could enable the packaging of the P68 genome in its head. Furthermore, the inner capsid proteins could function in determining the triangulation number of the P68 capsid during assembly, as was shown for Sid proteins of the *Enterobacteria* phage P4 of the P2/P4 system ^19^. The inner capsid proteins remain attached to the capsid after genome ejection, indicating that they do not participate in genome delivery (Fig. 1d).

### DNA is held inside the P68 head by interaction with one subunit of the portal complex

The portal complex of P68 is formed of twelve gp19 subunits (Fig. 3ab). It is 80 Å long along its twelvefold axis with the external diameters of the upper and lower parts of 140 and 100 Å, respectively (Fig. 3a). The structure of the P68 portal protein could be built except for residues 1-6 and 83-104 out of 327. According to the convention ^15^, it can be divided into three domains: the clip (residues 178-223), stem (6-41, 156-177, and 227-248), and wing (249-327) (Fig. 3a). Unlike the portal proteins of other phages, but similar to that of *Bacillus* phage phi29, the portal protein of P68 lacks a crown domain ^20–23^. The clip domain is composed of helix α4 and antiparallel strands β4 and β5. It forms part of the binding site for the lower collar complex and tail fibers. The stem domain of the portal protein is composed of a “tail fiber hook” (residues 6 – 11) and helices α1, α3, and α5 (Fig. 3a). The tail fiber hook enables the attachment of tail fibers to the portal complex, as discussed in detail below. Helices α1 and α3 of the stem domain form the outer surface of the portal complex that interacts with the capsid. The wing domain, which forms the part of the portal inside the phage head, consists of helices α2, α6-9 and strands β1-3. Helix α6 of the wing domain is inserted into the neighboring portal protein subunit and thus stabilizes the dodecamer structure of the portal complex (Fig. 3a).

**Fig. 3.**
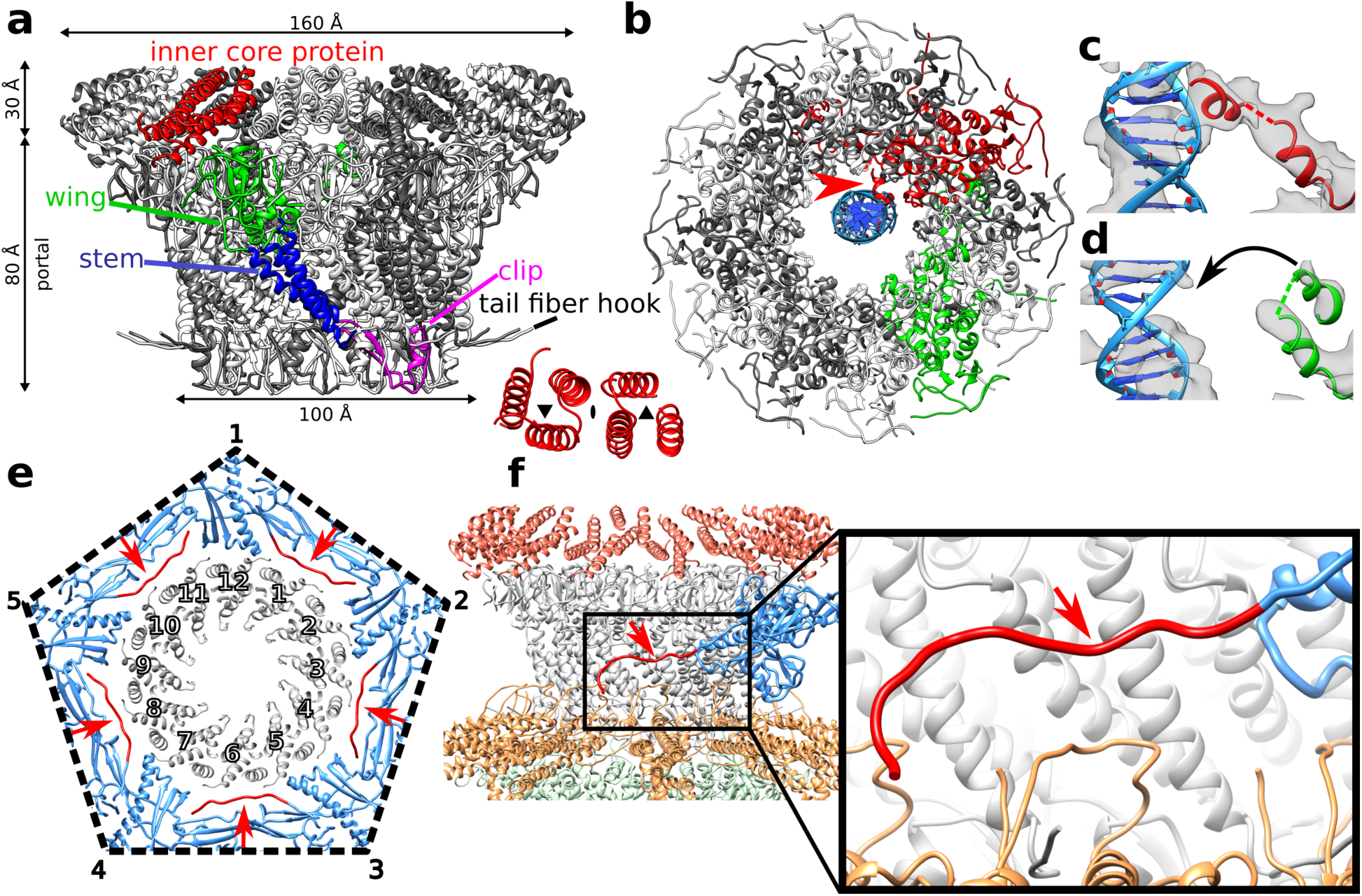
Structure of P68 portal complex and its interaction with capsid. Structure of P68 portal and inner core complexes (a). One of the portal proteins is colored according to domains: clip domain in yellow, wing in green, and stem in blue. Six inner core proteins associated with one portal protein subunit are highlighted in red. The inset shows the symmetry of the arrangement of the six inner core proteins. Asymmetric reconstruction of portal complex showing interactions of one of the portal proteins highlighted in red with DNA shown in blue (b). The interaction is indicated with a red arrow. One of the portal protein subunits that does not interact with the DNA is highlighted in green. Detail of interaction of helix α9 of portal protein with DNA (c). Cryo-EM density is shown as grey transparent surface. Structure of portal protein subunit that does not interact with DNA (d). Interface between portal complex and capsid (e). Portal proteins are shown in grey, capsid proteins in blue, and N-termini of capsid proteins that mediate interactions with the portal are shown in red and highlighted with red arrows. Side view of capsid-portal interactions (f). Single major capsid protein is shown in blue and its N-terminus in red, portal proteins are shown in grey, inner core proteins in red, tail fibers in orange and lower collar proteins in green. The inset shows detail of interactions between the N-terminal arm of the major capsid protein and stem domains of portal proteins.

Asymmetric reconstruction of the portal complex reveals a unique interaction of one of the portal proteins with the DNA positioned at the center of the portal channel (Fig. 3b-d). Helix α9 from the wing domain of the unique DNA-binding subunit binds to a side of the DNA helix (Fig. 3bc). It is the only observed interaction that may hold P68 DNA inside its head.

### Interface between capsid and portal complex

Asymmetric reconstruction of the entire P68 virion at a resolution of 4.7 Å enabled characterization of the interface between the capsid and portal complex (Fig. 3ef). Residues 1-42 from the N-terminal arm of the major capsid proteins adjacent to the portal are not structured. Residues 43-59 of the major capsid proteins wrap around the stem domains of portal proteins (Fig. 3f). If the major capsid proteins had the same structure as around the fivefold vertices occupied by capsid proteins, the N-terminal arms of the capsid proteins would clash with the portal and tail fibers (SFig. 6). It is likely that other tailed phages employ a similar mechanism for incorporating portal complexes into their capsids.

### Inner core proteins interact with the capsid and determine the attachment sites of head fibers

The surface of the portal complex facing towards the center of the P68 head is covered by seventy-two subunits of the inner core protein gp22 (Fig. 3a). Only residues 91-114, which form an α-helix, are resolved from each 147-residue long inner core protein. The six inner core proteins associated with each portal protein form two three-helix bundles related by a quasi-twofold rotational axis (Fig. 3a). Inner core proteins positioned closest to the capsid have additional structured residues that interact with axial domains of the adjacent major capsid proteins, and by modifying their conformations enable the attachment of head fibers to the capsid, as discussed below. Inner core proteins detach from the portal complex during the phage DNA release (Fig. 1c) and contain predicted pore-lining helices (SFig. 7) ^24^, indicating that they may enable the transport of the phage DNA across the bacterial cytoplasmic membrane.

### Head fibers can position P68 particles for genome delivery at cell surface

The P68 head is decorated with five trimers of head fibers gp14, which are attached to the hexamers of major capsid proteins adjacent to the tail vertex (Fig. 1a, 4ab). Low-resolution structures of head fibers are resolved in the asymmetric reconstruction of the P68 virion (Fig. 4a). Due to the mismatch of the fivefold symmetry of the head and twelvefold symmetry of the tail, only three head fibers are stabilized by interaction with the tail fibers (Fig. 4a). P68 head fibers can be divided into the N-terminal capsid-binding domain (residues 1-55), α-helical stalk (56-339), and receptor-binding domain (340-481), which is positioned at the level of the receptor binding domains of tail-fibers (Fig. 4a). A cryo-EM map of the P68 head enabled the building of the poly-alanine structure of 55 residues of the capsid-binding domain, which is composed of three beta-sheets and an α-helix (Fig. 4bc). Residues of the beta-sheets mediate the attachment of the head fiber to a hexamer of major capsid proteins (Fig. 4c). The alpha helices form a coiled coil that enables trimerization of the head fibers (Fig. 4bc). The selectivity of binding of the head fibers to the hexamers of major capsid proteins adjacent to the tail vertex is determined by interactions of the inner core proteins with the inner face of the capsid (Fig. 4de). Fifteen of the seventy-two inner core proteins present in the P68 head form structured “arms” that reach the axial domains of the closest hexamers of major capsid proteins (Fig. 4e). The interaction with the inner core proteins forces the side-chains of phe 259, from the axial domains of the major capsid proteins, to adopt a threefold symmetrical alternating “in and out” conformation (Fig. 4fg, SFig. 8a). In contrast, in hexamers of the major capsid proteins that do not interact with the inner core proteins, two phe 259 side chains point towards the center of the head and four side chains point away from the particle center (Fig. 4fh, SFig. 8b). In summary, the binding of the inner core proteins causes a change of symmetry of the six phe 259 side chains from twofold to threefold, and thus defines the attachment sites for the head fiber trimers.

**Fig. 4.**
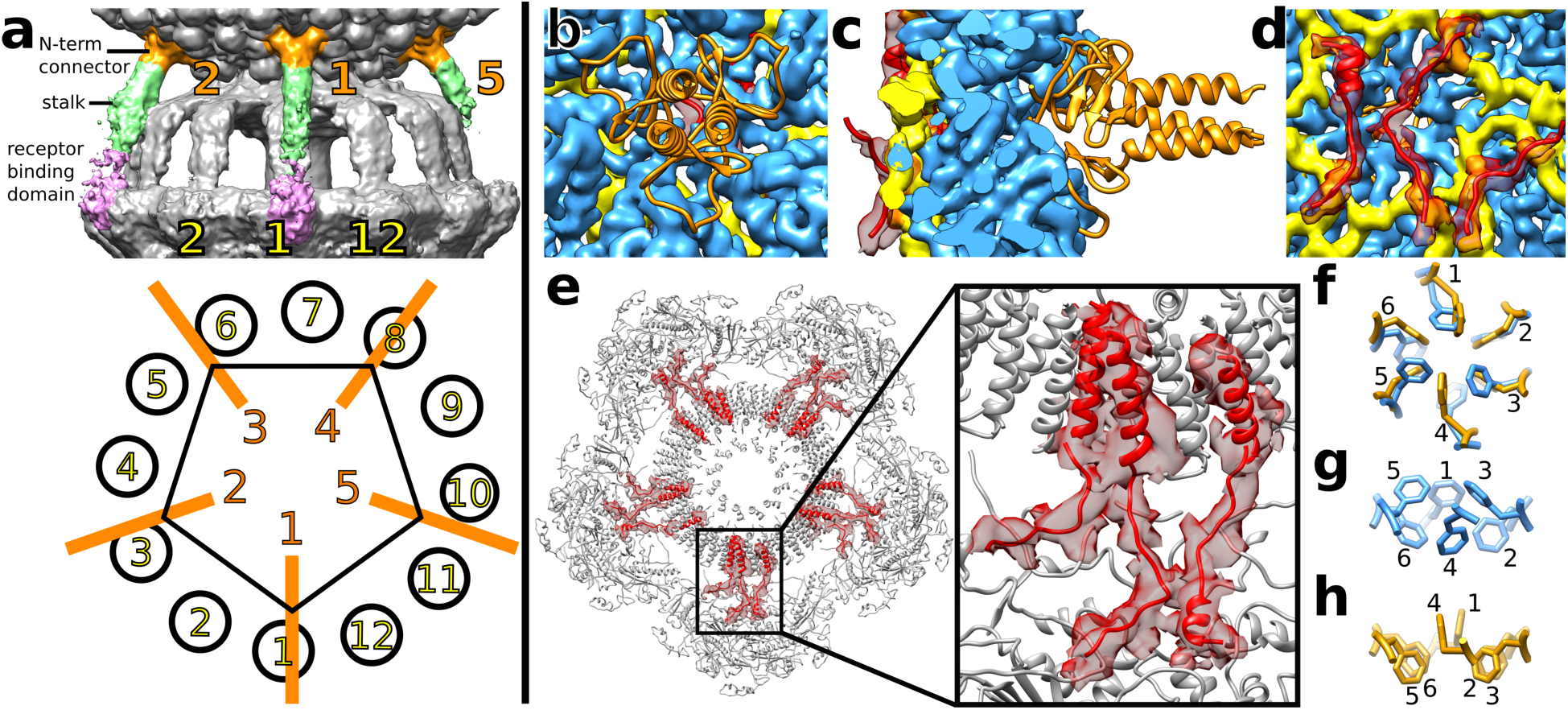
Head of P68 is decorated with five head fibers attached to hexamers of major capsid proteins located next to tail vertex. P68 head is decorated with five head fibers that extend towards tail fibers (a). The head fibers can be divided into the N-terminal connector shown in orange, stalk in green, and receptor binding domain in pink. Because of the mismatch of the fivefold symmetry of the head and twelvefold symmetry of the tail, only fibers 1, 2, and 4 are stabilized by interactions with tail fibers. N-terminal connector domains of head fibers (shown in cartoon representation in orange) are attached to hexamers of major capsid proteins (shown as blue density) (b-d). Cryo-EM density of inner capsid proteins is shown in yellow and arms of inner core proteins, which interact with major capsid proteins, are shown in cartoon representation in red. Cryo-EM density of inner core proteins is shown as semi-transparent red surface. External view of P68 head (b), section through capsid (c), and internal view of capsid (d). Section through P68 head perpendicular to tail axis at level of inner core complex (e). Inner core proteins that interact with major capsid proteins are highlighted in red. The electron density of the inner core proteins is shown as a red semi-transparent surface. Details of organization of phe 259 side chains around quasi-sixfold axis of hexamer of major capsid proteins (f-h). In hexamers that interact with inner core proteins, side-chains (in orange) are organized with threefold symmetry in alternating up and down conformations (f,g). In hexamers of capsid proteins that do not interact with inner core proteins, (blue) phe 259 side chains are organized with twofold symmetry with two side-chains pointing into capsid and four out (f,h).

Residues 340-477 of the head fiber are homologous to the receptor binding proteins of lactococcal phages TP901-1, P2, and bIL170 (Table. S3) ^25–27^. The putative receptor binding domains of P68 head fibers are positioned next to the receptor binding domains of tail fibers (Fig. 4a). Therefore, the binding of head fibers to receptors can position P68 with its tail orthogonal to the cell surface for genome delivery. This is supported by the broader host range of phage P68 in comparison to the closely related phage 44AHJD, which lacks the gene for the head fiber ^13,28^. In contrast, the head fibers of previously structurally characterized phages point in all directions and are thought to function in the reversible attachment of phages to cells in random orientations.

### Lower collar complex of P68 tail

The lower collar complex is attached to the portal complex and forms the central part of the P68 tail (Fig. 1ab, 5a). The dodecamer of lower collar proteins (gp18) has the shape of a mushroom with a head diameter of 162 Å and total length of 146 Å (Fig. 5a). It contains an axial channel that is continuous with that of the portal complex (Fig. 1b). The structure of the lower collar protein could be built except for residues 1 and 154-186 out of 251. It can be divided into three parts: the curly domain (residues 3–116 and 222–251), tube domain (116–154 and 184 – 222), and knob connector loop (154 – 184) (Fig. 5b). The curly domain, formed by six α-helices, mediates the attachment of the lower collar complex to the portal complex and tail fibers. The tube domain is composed of two antiparallel β-strands (Fig. 5b). Twelve tube domains form a β-barrel with twenty-four β-strands, which is 108 Å long (Fig. 5a). The knob connector loops enable the attachment of the tail knob complex, with sixfold symmetry, to the lower collar complex.

**Fig. 5.**
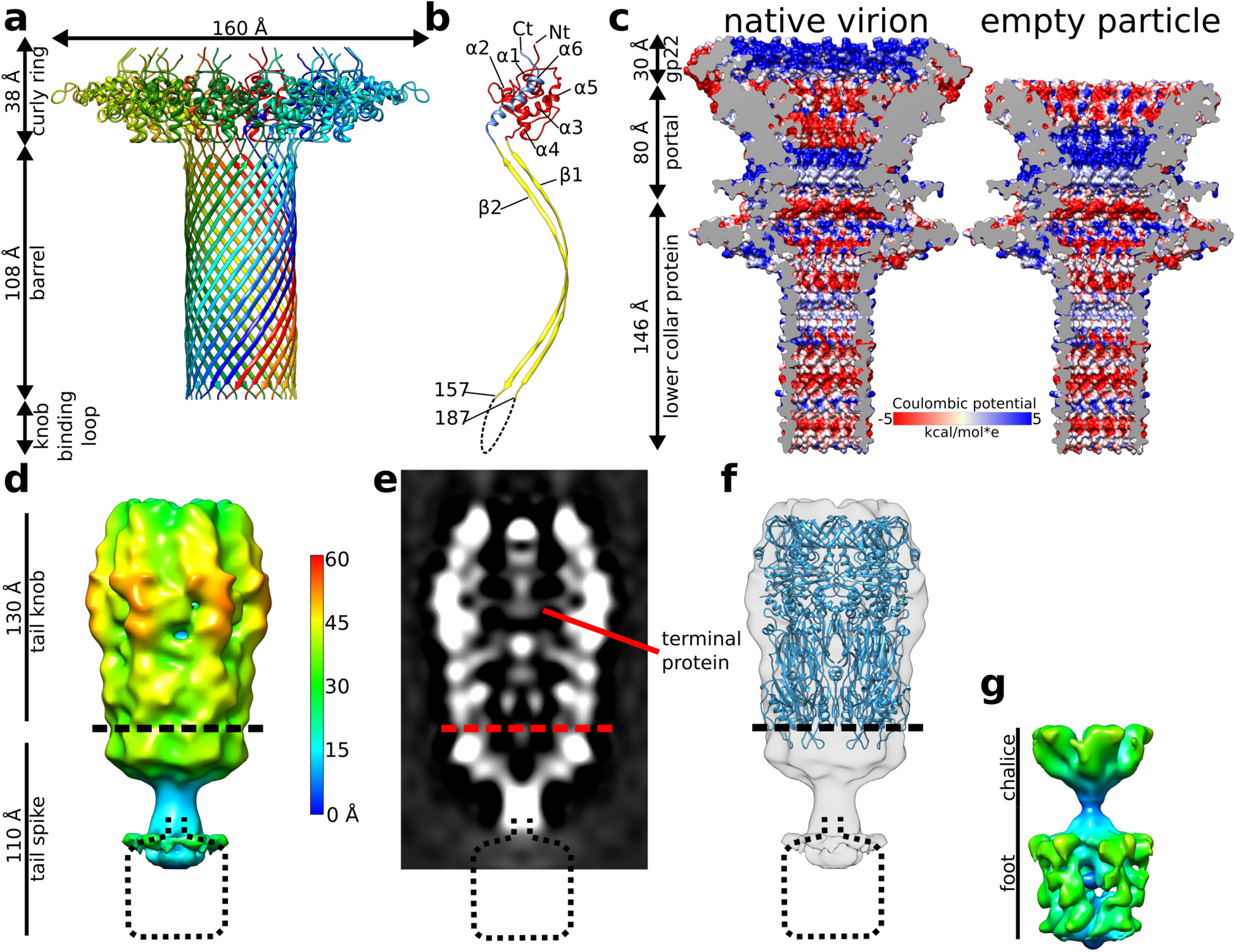
Structure of P68 tail. Structure of lower collar complex (a) with individual subunits distinguished by rainbow-coloring. Division of lower collar protein into domains (b). The curly ring domain is shown in red and blue, barrel domain in yellow, and the knob binding loop with unknown structure is indicated by the dashed line. Surface charge distribution in inner core, portal, and lower collar complexes of full and empty P68 particles (c). Sixfold-symmetrized reconstruction of P68 tail knob and tail spike complexes (d). The surface of the cryo-EM map is radially colored based on the distance from the sixfold axis of the complex. Distribution of electron density in central section of tail knob and tail spike complexes (e). Fit of structure of tail knob of phage p22 into P68 reconstruction (f). Structure of tail spike with imposed fivefold symmetry shows its chalice and foot domains (g).

Portal and lower collar complexes of P68 form a channel with a total length of 270 Å (Fig. 5c). The inner diameter of the channel varies from 30 Å to 55 Å. The surface charge distribution inside the channel is mostly negative, but it is interrupted by neutral and positively charged layers (Fig. 5c).

### Tail knob and tail spike

The tail of P68 continues beyond the lower collar protein by tail knob gp13 and tail spike gp11 (Fig. 1ab). The two complexes were reconstructed to resolutions of 11 Å and 7 Å, respectively, indicating that they are more flexible than the parts of the tail near the phage head (Fig. 5d-g). This flexibility may be required to allow the putative cell wall-degrading enzymes located in the tail spike to cleave a pore in the bacterial cell wall to enable genome delivery.

The previously determined crystal structure of the tail knob of *Streptococcus* phage C1 fits into the reconstruction of the corresponding part of the P68 tail with a correlation coefficient of 0.65 (Fig. 5f) (Table S4) ^29^. The length of the P68 tail knob is 130 Å along the tail axis. It has an outer diameter of 80 Å and inner tube diameter of 40 Å. The channel is continuous with that of the lower collar protein (Fig. 1b). The tail of native P68 contains a tubular density that may belong to a terminal protein (Fig. 1b, 5e), which is covalently linked to the end of P68 DNA ^13^.

An asymmetric reconstruction of the tail spike provides evidence that it has fivefold symmetry (SFig. 9). Fivefold symmetrized, localized reconstruction of the tail spike shows that it is 110 Å long and has a maximum diameter of 70 Å (Fig. 5g). It can be divided into the chalice,which mediates attachment to the tail knob, and distal lysis domain (Fig. 5g). Sequence comparisons indicate that the tail spike of P68 is homologous to the PlyCb lysin from phage C1 (Table S5) ^30^. However, the structure of PlyCb does not fit into the reconstruction of the P68 tail spike ^31^. Other proteins with peptidoglycan degradation activities such as the amidase from *S. aureus*, peptidases from *Staphylococcus saprophyticus*, and endolysin from staphylococcal phage K are homologous to the last 130 amino acids of the P68 tail spike protein (SFig. 9). This indicates that the tail spike proteins of P68 degrade the bacterial cell wall to enable access of the phage to the cytoplasmic membrane.

### Tail fibers

The tail fibers of P68 form a skirt-like structure around the tail (Fig. 1a, Fig 6a). Each tail fiber is a trimer of 647-residue-long gp17 subunits (Fig. 6b). The tail fiber can be divided into the N-terminal stem domain (residues 1-145), platform (151-445), and C-terminal tower (446-647) (Fig. 6b).

**Fig. 6.**
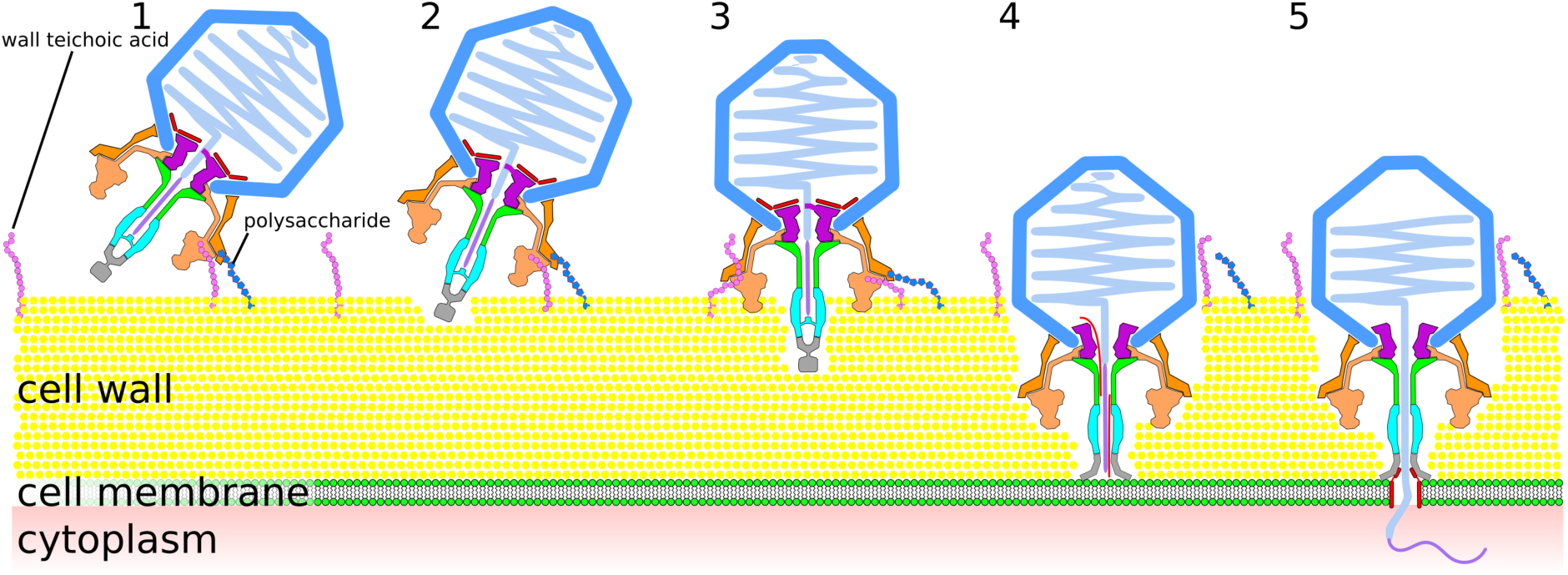
Structure of P68 tail fibers. Tail fibers of P68 form “skirt” around tail tube (a). Tail fibers are shown in orange, however, individual subunits of two tail fibers are distinguished in red, blue, and orange. Portal proteins are shown in magenta, inner core proteins in dark grey, and lower collar proteins in light grey. The inset shows details of interactions of tail fibers with each other and their attachment to the portal complex. Structure of P68 tail fiber in cartoon representation and its division into domains (b). The inset shows a cartoon representation of the platform domain of the P68 tail fiber rainbow colored from N-terminus in blue to C-terminus in red.

Cryo-EM reconstruction of the P68 tail enabled the building of the structure of the stem domain, which can be further sub-divided into a connector (residues 1-45), shoulder (46-80), hinge (81-115), and arm (116-145) (Fig. 6b). Because of the asymmetric shape of the tail fiber, the three constituent subunits (A, B, and C) differ in structure from each other (Fig. 6b). Functionally important differences are found in the connector regions that mediate the attachment of the tail fiber to the portal and lower collar complexes (Fig. 6a). The connector domain of subunit A and shoulder from subunit C form a noose-like structure that encircles the N-terminal tail-fiber hook of the portal protein (Fig. 6a). The connector of subunit B binds to the shoulder domains of subunits A and B from the tail fiber positioned counterclockwise when looking at the tail from the direction of the head (Fig. 6a). The first structured residue of the connector of subunit C (thr 24) is located between the clamp domain of subunit B from the tail fiber positioned clockwise and shoulder domain of subunit C and the clamp domain of subunit A positioned counterclockwise (Fig. 6a). Thus the N-terminus of C subunit mediates interactions between tail fibers that are one position removed from each other.

The shoulder domain of the tail fiber is straight until the hinge domain, which introduces a turn of 110° (Fig. 6b). The hinge of subunit C is formed by two α-helices connected by a short loop, which allows the chain to bend and pass under subunits A and B (Fig. 6b). After the hinge, the three subunits form a straight coiled-coil arm (Fig. 6b).

The cryo-EM reconstruction of the P68 tail is complemented by the crystal structure of the tail-fiber protein determined to a resolution of 2.0 Å (Table S6). Although the full-length tail fiber protein was used for crystallization, only the platform and tower domains (residues 139-647) were resolved (Table S6). The combination of cryo-EM and X-ray results allowed construction of the complete tail fiber.

The platform domain of the P68 tail fiber has a five-bladed β-propeller fold (Fig. 6b). Each of the blades contains four anti-parallel β-strands. The domain is cyclically enclosed, since the first N-terminal β-strand of the domain is part of the same blade as the last three C-terminal β-strands (Fig. 6b). It has been shown that the platform domains of various phages, including phi11, PRD1, and PhiKZ, contain receptor-binding sites ^32–34^. The platform of the P68 tail fiber is similar in structure to that of staphylococcal phage phi11 from the family *Siphoviridae*, with an RMSD of the corresponding Cα atoms of 1.10 Å. The sequence identity of the two proteins is 24%. There are differences in the receptor-binding sites within the platform domains of the two phages that may reflect their different receptor requirements (SFig. 10). Whereas P68 binds to wall teichoic acid glycosylated with β-O-N-acetyl-glucosamine, phi11 can attach to both β-O-N-acetyl-glucosamine and α-O-N-acetyl-glucosamine ^2^.

The tower domain of the P68 tail fiber is composed of two sub-domains (residues 454 – 555 and 556 – 645), which are structurally similar to each other (Fig. 6b). Each sub-domain is formed of a four-stranded antiparallel beta-sheet connected by loops and two short helices (Fig. 6b). The beta-sheets are positioned close to the threefold axis of the fiber, whereas the loops and helices are exposed at the surface. The sub-domains are homologous to putative major teichoic acid biosynthesis protein C, muramidases, and receptor binding fibers of R-type pyocin (Table S7) ^35^. Thus, the tower region may be involved in binding to the cell wall or peptidoglycan digestion. Compared to the tail fiber of phage phi11, the platform and tower domains of P68 exhibit domain swapping within the trimer of the tail fiber (SFig. 11) ^32^.

The cryo-EM structure of P68 tail shows interactions of residues 229-271 of platform domain of subunit B with residues 349-386 of chain A and 180-268 of chain B of platform domains from the neighboring tail fiber (Fig. 1ab, Fig. 6a). The interface has a buried surface area of 1,100 Å^2^ and it is likely that it stabilizes the “skirt” structure of P68 tail fibers. The flexibility of the hinge region of tail fibers was proposed to facilitate receptor binding in other phages ^32^. In contrast, in P68 the structure of tail fibers appears to be rigid.

### Changes in P68 particles associated with genome release

The genome release of P68 is connected to the disruption of the unique contact of the portal protein subunit with the DNA, loss of the ordering of most of the wing domains of the portal proteins and of the inner core proteins (Fig. 1b-d, SFig. 12). Unlike phage phi29, P68 particles do not bind to liposomes at low pH (SFig. 13) ^36^, instead they aggregate with each other through their tails (SFig. 13). However, liposomes in the mixture with P68 became distorted (SFig. 13). It is possible that the ejected inner core proteins, which contain predicted pore-lining helices (SFig. 7), interfered with the liposome integrity.

The heads of P68 particles in the process of genome release contain shells of packaged DNA that are spaced 26-30 Å apart, whereas in the full virions the DNA spacing is 20 Å (Fig. 1bc, SFig. 14). The resolved structure of the layers of dsDNA in the P68 genome release intermediate indicates that all the particles released similar amounts of DNA and provides evidence of a gradual relaxation of the DNA packing during the genome release.

Only 2% of P68 genome release intermediates and empty particles retained their tail knobs and tail spikes *in vitro* (SFig. 1). However, the tail knobs and tail spikes of the complete empty particles contain central channels (Fig. 1d), indicating that the complexes may remain attached to P68 virions during genome release *in vivo*.

### Mechanism of P68 genome delivery

P68 virions bind to the *S. aureus* cell either by head or tail fibers (Fig. 7). After the attachment, tail spike proteins degrade the cell wall, which allows the phage to position itself with its tail axis perpendicular to the cell surface (Fig. 7). Further cell wall digestion enables the tip of the P68 tail to reach the *S. aureus* cytoplasmic membrane. The signal triggering P68 genome release is unknown, however, it may be the binding of the tail spike to a receptor in the membrane, exposure of the tail spike to the hydrophobic environment of the membrane, or a sensing of the trans-membrane potential ^1^. Subsequently, conformational changes of the P68 portal enable ejection of the inner core proteins and DNA through the tail channel. The inner core proteins may form a pore in the bacterial membrane for delivering phage DNA into the bacterial cytoplasm (Fig. 7).

**Fig. 7.**
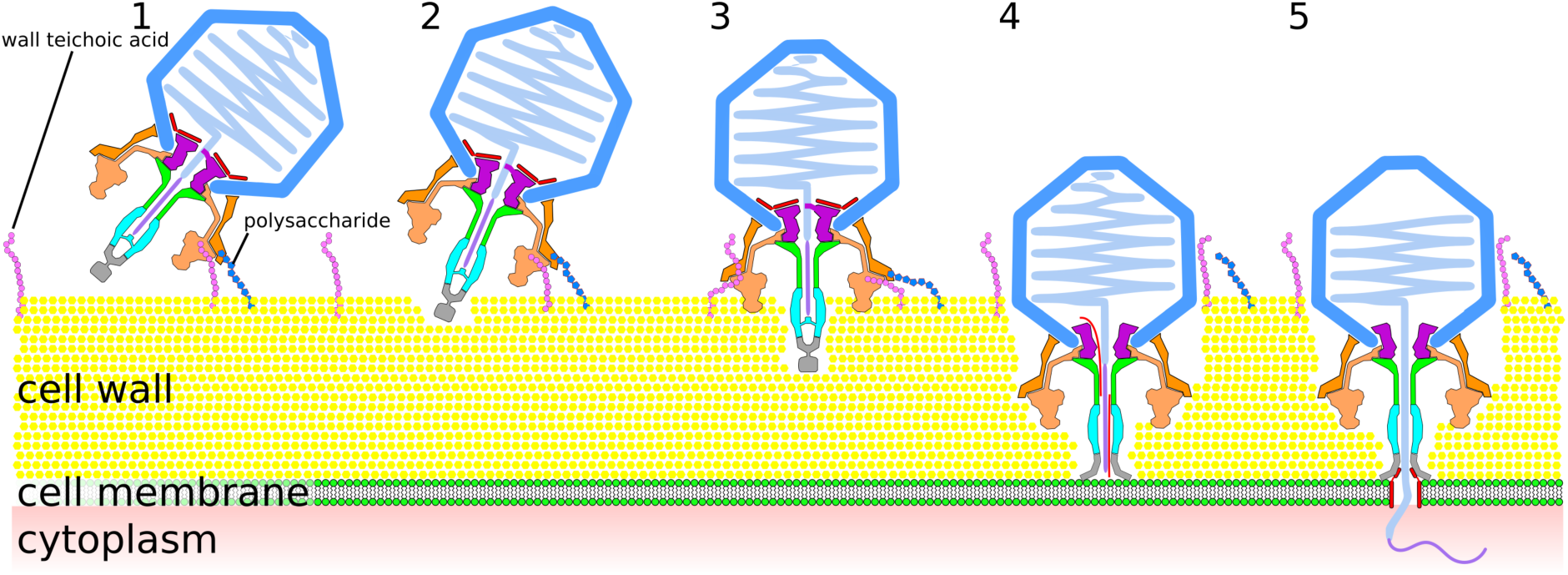
Mechanism of P68 genome delivery into *S. aureus* cell. P68 virion attaches to cell surface by head or tail fibers (1). This attachment allows enzymes from tail spike to cleave bacterial cell wall (2). This degradation of *S. aureus* cell wall enables P68 to bind with its tail axis perpendicular to cell surface (3). Further cell wall digestion allows tip of P68 tail to reach cytoplasmic membrane, which triggers release of inner core proteins and DNA (4). Inner core proteins form channel in membrane for ejection of phage DNA into bacterial cytoplasm (5).

## Acknowledgements

We acknowledge the CEITEC Core Facilities Cryo-Electron Microscopy and Tomography, Proteomics, and Biomolecular Interactions, supported by the CIISB project LM2015043 funded by the MEYS of the Czech Republic. X-ray data were collected at synchrotron ‘Soleil’ beamline Proxima 1. This research was carried out under the project CEITEC 2020 (LQ1601), with financial support from the MEYS of the Czech Republic under National Sustainability Program II. This work was supported by IT4I project (CZ.1.05/1.1.00/02.0070), funded by the European Regional Development Fund and the national budget of the Czech Republic via the RDI-OP, as well as the MEYS via the Grant (LM2011033). The research leading to these results has received funding from the Ministry of Health of the Czech Republic grant 16-29916A to RP, Grant Agency of the Czech Republic grants 15-21631Y and 18-17810S and from EMBO installation grant 3041 to PP.

